# A distinct isoform of Msp300 (nesprin) organizes the perinuclear microtubule organizing center in adipocytes

**DOI:** 10.1101/2024.06.28.601268

**Authors:** Garret M. Morton, Maria Pilar Toledo, Chunfeng Zheng, Yiming Zheng, Timothy L. Megraw

## Abstract

In many cell types, disparate non-centrosomal microtubule-organizing centers (ncMTOCs) replace functional centrosomes and serve the unique needs of the cell types in which they are formed. In Drosophila fat body cells (adipocytes), an ncMTOC is organized on the nuclear surface. This perinuclear ncMTOC is anchored by Msp300, encoded by one of two nesprin-encoding genes in Drosophila. Msp300 and the spectraplakin short stop (shot) are co-dependent for localization to the nuclear envelope to generate the ncMTOC where they recruit the microtubule minus-end stabilizer Patronin (CAMSAP). The *Msp300* gene is complex, encoding at least eleven isoforms. Here we show that two Msp300 isoforms, Msp300-PE and -PG, are required and only one, Msp300-PE, appears sufficient for generation of the ncMTOC. Loss of Msp300-PE and -PG results in severe loss of localization of shot and Patronin, disruption of the MT array, nuclear mispositioning and loss of endosomal trafficking. Furthermore, upon loss of Msp300-PE and -PG, other isoforms are retained at the nuclear surface despite the loss of nuclear positioning and MT organization, indicating that they are not sufficient to generate the ncMTOC. Msp300-PE has an unusual domain structure including a lack of a KASH domain and very few spectrin repeats and appears therefore to have a highly derived function to generate an ncMTOC on the nuclear surface.

## Introduction

The <u>li</u>nker of <u>n</u>ucleoskeleton and <u>c</u>ytoskeleton (LINC) complex spans the nuclear envelope and provides a physical connection between the nuclear interior with the cytoplasm and plasma membrane [1, 2]. The essential roles of the LINC complex in cell function are underscored by inherited diseases associated with mutations in LINC complex protein genes [3, 4] including muscular dystrophies [5, 6], cardiomyopathies [7, 8], cerebellar ataxia [9, 10], and arthrogryposis [11]. The LINC complex is integral to cellular processes such as DNA damage response [12, 13], meiotic chromosome movement [14, 15], mechanosensation [1], and nuclear positioning and anchoring [3, 16–19]. Comprising the LINC complex are SUN (<u>S</u>ad1/<u>UN</u>C-84 homology) and KASH (<u>K</u>larsicht/<u>A</u>NC-1/<u>S</u>yne <u>h</u>omology) proteins. SUN proteins reside at the inner nuclear membrane and contain an N-terminus that extends into the nucleoplasm, where it interacts with lamins of the nucleoskeleton and other nucleoplasm proteins, while the C-terminal SUN domain forms contacts with KASH proteins in the perinuclear space. The KASH proteins reside on the outer nuclear membrane, associate with cytoskeletal components, and are characterized by a C-terminal KASH domain that associates with the SUN domain of the SUN proteins within the nuclear intermembrane space. In the absence of a SUN domain protein interaction in the intermembrane space, the KASH protein does not localize to the outer nuclear membrane [1, 20].

In *Drosophila*, the LINC complex components include two KASH proteins, Klarsicht (Klar) and <u>M</u>uscle-<u>s</u>pecific <u>p</u>rotein <u>300</u> kDa (Msp300), and one primary SUN protein, Klaroid (Koi), while a second SUN protein, Spag4, is expressed exclusively in the testes. The Drosophila LINC proteins are responsible for the proper positioning of muscle myonuclei [21], similar to their roles in vertebrate muscle [6, 22–24]. Msp300, which was first identified in the muscle [25], promotes myonuclear spacing [26]. These findings implicate nuclear mispositioning in the pathogenesis of diseases such as Emery-Dreifuss muscular dystrophy (EDMD) [26, 27]. As seen with other <u>n</u>uclear <u>e</u>nvelope <u>sp</u>ectrin <u>r</u>epeat prote<u>in</u>s (nesprins), Msp300 contains calponin homology (CH) domains, numerous spectrin repeats (SR), and a C-terminal KASH domain. In addition, *Msp300* encodes at least eleven isoforms of various molecular weights expressed from a complex locus (Figure 1). Furthermore, subsets of these isoforms have been shown to exhibit unique functions. In the muscle, isoforms containing the CH domains but lacking the KASH domain were found to contribute to Z-line organization and for optimal muscle function, while isoforms lacking the CH domains but containing the KASH domain were shown to be responsible for maintaining myonuclear positioning [28].

**Figure 1.**
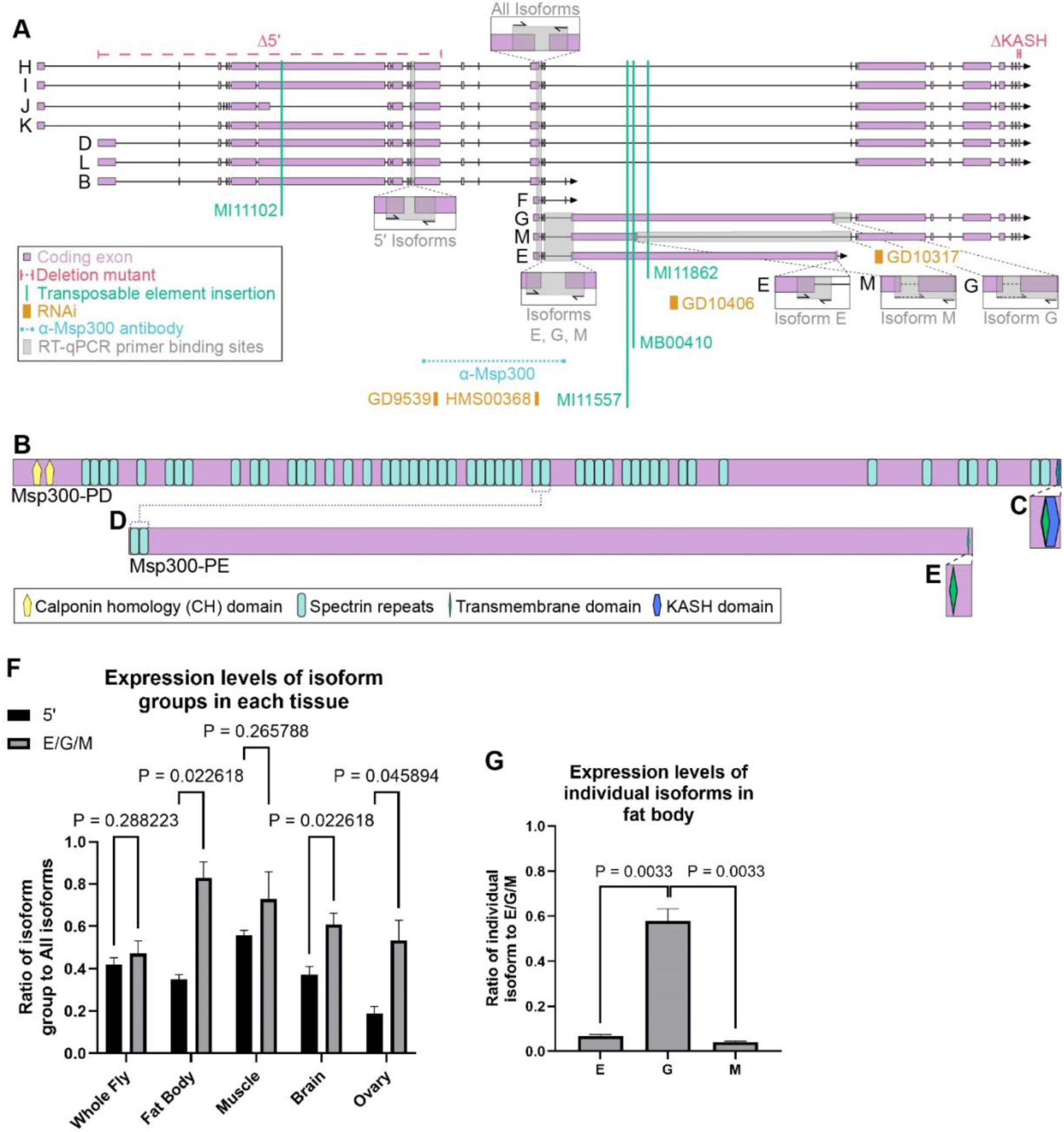
*Msp300* gene isoforms and genetic analysis features. (A) Schematic of the isoforms expressed from *Msp300*. Deletion mutants (red), transposable element insertion mutations (green), and RNAi (orange) are indicated. The isoforms and gene regions affected by these mutations or RNAi elements are depicted. The epitope recognized by the α-Msp300 antibody (blue) is shown. Binding sites of the primers for RT-qPCR are represented by magnified views. (B-E) Schematic representations of Msp300-PD, a ‘conventional’ Nesprin, and Msp300-PE. Nesprins and most Msp300 isoforms such as PD (B) characteristically contain calponin homology domains, numerous spectrin repeats, and a KASH domain associated with a transmembrane domain (C). The atypical Msp300-PE (D) lacks calponin homology domains and contains only two spectrin repeats. The majority of Msp300-PE is comprised of two types of tandem repeats. Isoform PE also contains a transmembrane domain but lacks a KASH domain (E). (F) Bar graph comparing the mRNA expression levels of isoform groups relative to all isoforms in different tissues from RT-qPCR data from indicated tissues. RNA extractions were performed in triplicate and each RT-qPCR reaction was run in triplicate. Expression levels were normalized to GPDH1 and α-Tubulin reference genes. (G) Expression levels of isoforms E, G, and M relative to the expression level of all three isoforms combined in the fat body. The data are depicted as the mean ± SD. Statistical significance was determined using a two-tailed Student’s *t*-test with Welch’s correction (*n* = 3). See S4 Data for source data.

*Drosophila* fat body cells, which are functionally comparable to mammalian adipocytes and liver cells, are characterized by a nucleus positioned at the center of the cell and MTs organized at the nuclear surface that radiate outwards and connect the nucleus to the plasma membrane [29, 30]. The perinuclear fat body ncMTOC controls critical cellular functions, including microtubule organization, nuclear positioning, and endocytic vesicle trafficking [29]. Two general classes of MTs are organized at the perinuclear ncMTOC, circumferential MTs surround the nucleus, while radial MTs extend from the nucleus into the cytoplasm. The control of MT assembly at the fat body perinuclear ncMTOC has unique molecular features. Assembly of MTs at this ncMTOC requires Patronin, Ninein, and msps (ortholog of ch-TOG), but does not require γ-tubulin [29]. Patronin (CAMSAP ortholog) is a MT stabilizer involved at many ncMTOCs [31–37], Ninein is a MT anchor that associates with dynein light intermediate chain [38–40] and with ensconsin/MAP7 and cooperates with ensconsin in MT assembly [41–43], and msps/ch-TOG is a MT polymerase that associates with Patronin [29, 44]. Notably, Msp300 is the only LINC complex protein that establishes the ncMTOC, while Klar and Koi are not required [29], suggesting a novel and unconventional role for Msp300 at the fat body ncMTOC. Here, we have further explored the complex biology of Msp300 isoforms and identified one particular and distinct isoform of Msp300 that is responsible for organization of the perinuclear ncMTOC in the fat body. By utilizing deletion mutants, transposable element insertion mutations, and RNAi, we show that one unconventional isoform, Msp300-PE, appears necessary and sufficient for establishment of the ncMTOC on the nuclear surface.

## Results

### The *Msp300* locus is complex

The *Msp300* gene is complex, encoding at least 11 isoforms that range in size from 47 to 1,500 kDa (Fig 1A). Based on the curated data on FlyBase, these variants arise from alternative promoters, mRNA splice sites, and transcriptional terminators [45]. The existence of the different isoforms is supported by RNA-seq data of exon-exon junctions and the isolation of partial cDNA clones for all isoforms [46–48]. In addition, tissue-specific expression data of the isoforms indicates that all isoforms are expressed in the fat body [49]. Typically, nesprins consist of many spectrin repeats (SRs), a pair of calponin homology (CH) domains, and a KASH domain [50]. From available expression data, isoforms B, E, and F lack the KASH domain, and isoforms E, F, G, and M lack CH domains. The KASH domain consists of a transmembrane helix and a short C-terminal region that resides in the nuclear intermembrane space [51]. The human giant nesprin isoforms, Nesprin-1 (1.0 MDa) and Nesprin-2 (0.8 MDa), contain 74 and 56 SRs, respectively [52, 53]. Similarly, one of the largest Msp300 isoforms, Msp300-PD (1.41 MDa), contains 52 SRs, as well as 2 CH domains and a KASH domain (Fig 1B-C) [54]. Msp300-PG (1.5 MDa) contains no CH domains, 21 SRs, and a KASH domain. Msp300-PE (1.06 MDa) is an atypical nesprin with no CH domains, only 2 SRs, and no KASH domain (Fig 1D). Instead of a KASH domain, Msp300-PE is predicted to have a transmembrane domain near its C-terminus (Fig 1E), followed by a 35 aa sequence.

### Msp300 isoforms E/G/M are expressed more highly in the fat body

We measured the mRNA expression levels of groups of Msp300 isoforms in different tissues, including whole body, brain, flight muscle, ovary and fat body, by reverse transcription quantitative PCR (RT-qPCR). We assessed the expression of three sets of isoform mRNAs: all isoforms, those expressed from the two most 5′ promoters, and the E/G/M group of isoforms (Fig 1A). These data show that *Msp300* is expressed in all tissue types tested (Fig 1F). In the fat body, E/G/M are more highly expressed than the 5′ isoforms. Of the E/G/M group of isoforms in the fat body, isoform G was expressed at the highest level (Fig 1G). In addition, the existence of isoform E is validated by the RT-qPCR expression data.

### Knockdown of Msp300 isoforms E and G impair the ncMTOC

To further assess which protein isoforms of *Msp300* are expressed and localized to the nuclear envelope in fat body cells, we used RNAi to knock down expression of isoform groups and stained with a polyclonal Msp300 antibody that targets all isoforms (Fig 1A) to detect remaining isoforms at the nuclear envelope. We generated tissue mosaics of RNAi knockdown using the coinFLP system [55] (Fig 2). With coinFLP, we obtained mosaic tissues where RNAi knockdown cells were generated efficiently at a reliable ratio with control cells, affording an internally-controlled framework to assess RNAi phenotypes. The RNAi knockdown clones are differentiated from control cells by co-expression of nuclear-targeted mCherry (mCherry-nls) in the clones. As an additional control, wild-type (WT) clones were generated in the *w^1118^* background (Fig 2A, F).

**Figure 2.**
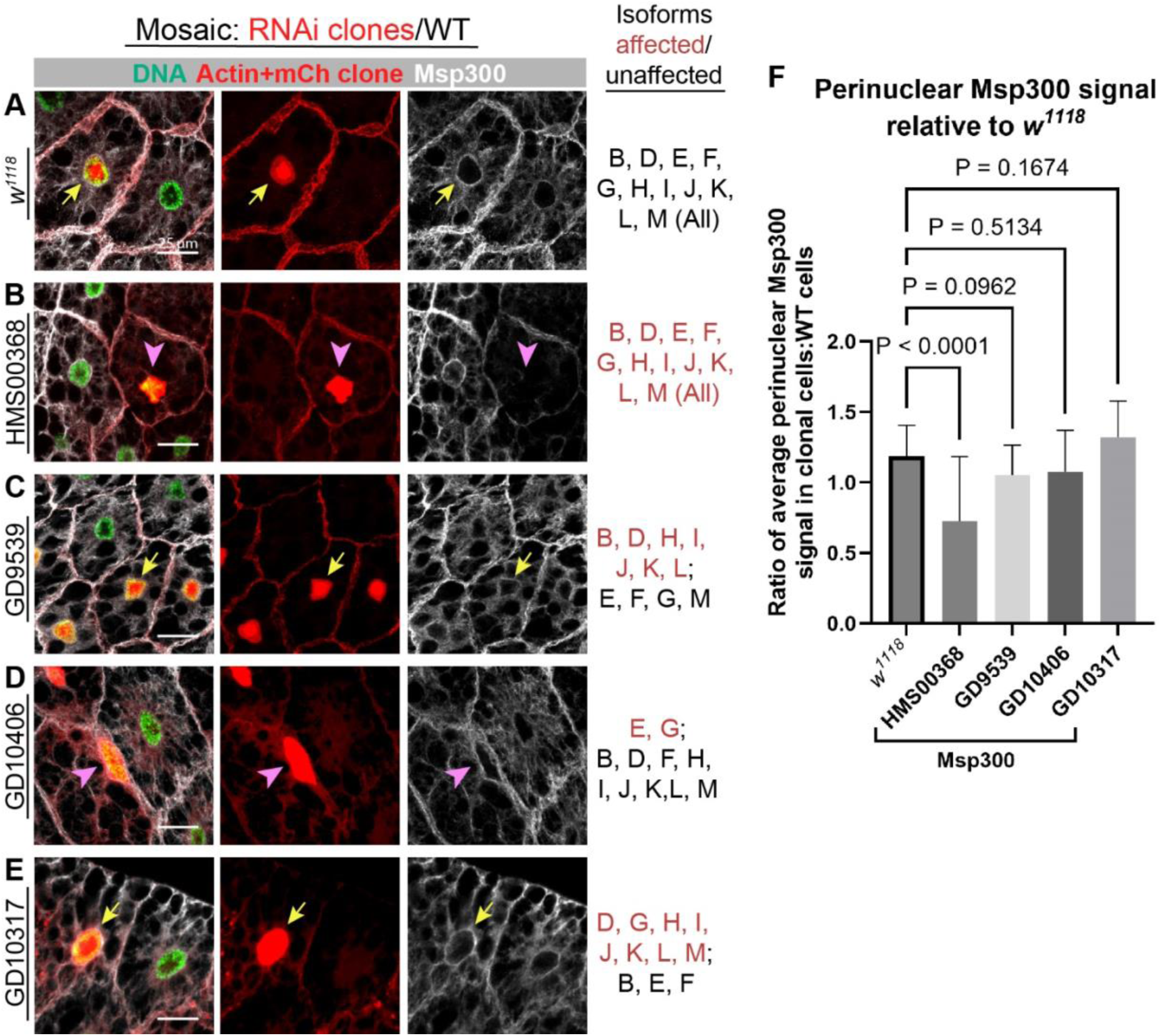
Multiple Msp300 isoforms localize to the nuclear surface. Immunostaining of fat bodies for Msp300 in *w^1118^* (A) or following RNAi knockdown by the HMS00368 (B), GD9539 (C), GD10406 (D), or GD10317 (E) RNAi lines. Images shown are representative of the phenotype associated with each line. Mosaic knockdown was achieved using the coinFLP system. Clonal RNAi knockdown cells express nuclear mCherry (red). Nuclei were stained with DAPI and pseudo-colored green, actin (cell membrane) stained red, and Msp300 white. Isoforms targeted by each RNAi line are indicated. Clones characterized by nuclear mispositioning are denoted by a pink arrowhead, while clones with properly positioned nuclei are denoted by a yellow arrow. (F) Quantitation of perinuclear Msp300 localization. Data are represented as the mean ± SD. A one-way ANOVA with Dunnett’s T3 test was used (24 ≤ *n* ≤ 30). Source data is found in S4 Data.

When all Msp300 isoforms were knocked down (with HMS00368 [56]), the perinuclear Msp300 signal was significantly reduced compared to the control cells, as expected (Fig 2B, F). Consequentially, the nuclei were mispositioned as previously shown for *Msp300* knockdown due to the loss of microtubule organization at the nuclear surface [29]. In contrast, targeting of the 5′ isoforms (with GD9539 [57]) did not diminish Msp300 localization to the nuclear surface (Fig 2C, F). Knockdown of isoforms E and G (with GD10406 [57]) results in nuclear mispositioning; however, the Msp300 signal at the nuclear surface was not significantly reduced (Fig 2D, F). The remaining Msp300 signal at the nuclear envelope is likely contributed by the other isoforms not targeted by RNAi. Meanwhile, knockdown of all KASH-containing isoforms (with GD10317 [57]), which includes G, does not significantly reduce perinuclear Msp300 (Fig 2E, F). The persistence of Msp300 signal at nuclei following knockdown of each set of transcripts is consistent with the RT-qPCR data showing that most Msp300 isoforms are expressed in the fat body. Importantly, the loss of nuclear positioning upon knockdown of the E/G isoforms, while retaining Msp300 signal from other isoforms, indicates that Msp300-PE and -PG are uniquely required for nuclear positioning, likely through the regulation of MT organization. The retention of Msp300 signal despite loss of nuclear positioning indicates that other isoforms cannot compensate and lack the function to organize the ncMTOC and that this role is unique to the Msp300-PE,-PG isoforms.

### Msp300 isoform E is sufficient for nuclear positioning and microtubule organization, with probable contribution from Isoform G

To determine which isoforms are required to maintain nuclear positioning and MT organization, we examined phenotypes for a variety of mutant lines in addition to the RNAi lines used in Figure 2. We generated mosaic mutant clones using the FLP-FRT recombination system [58], in which control cells express nuclear RFP while mutant cells do not (Fig 3). First, we examined two deletion mutants that impact groups of isoforms: one that deletes the 5′ end of the gene (*Msp300^Δ5’^*) [59], disrupting isoforms B, D, H, I, J, K, and L, and a second deletion mutant that deletes a segment encoding the KASH domain (*Msp300^ΔKASH^*) [60] (see Fig 1A). In both mutants, nuclear positioning and MT organization were unaffected, indicating that the 5′ isoforms and also the KASH domain are not required for generating the ncMTOC (Fig 3A-B, G-H). The lack of requirement for the KASH domain indicates that the LINC complex involved in generating the ncMTOC is atypical.

**Figure 3.**
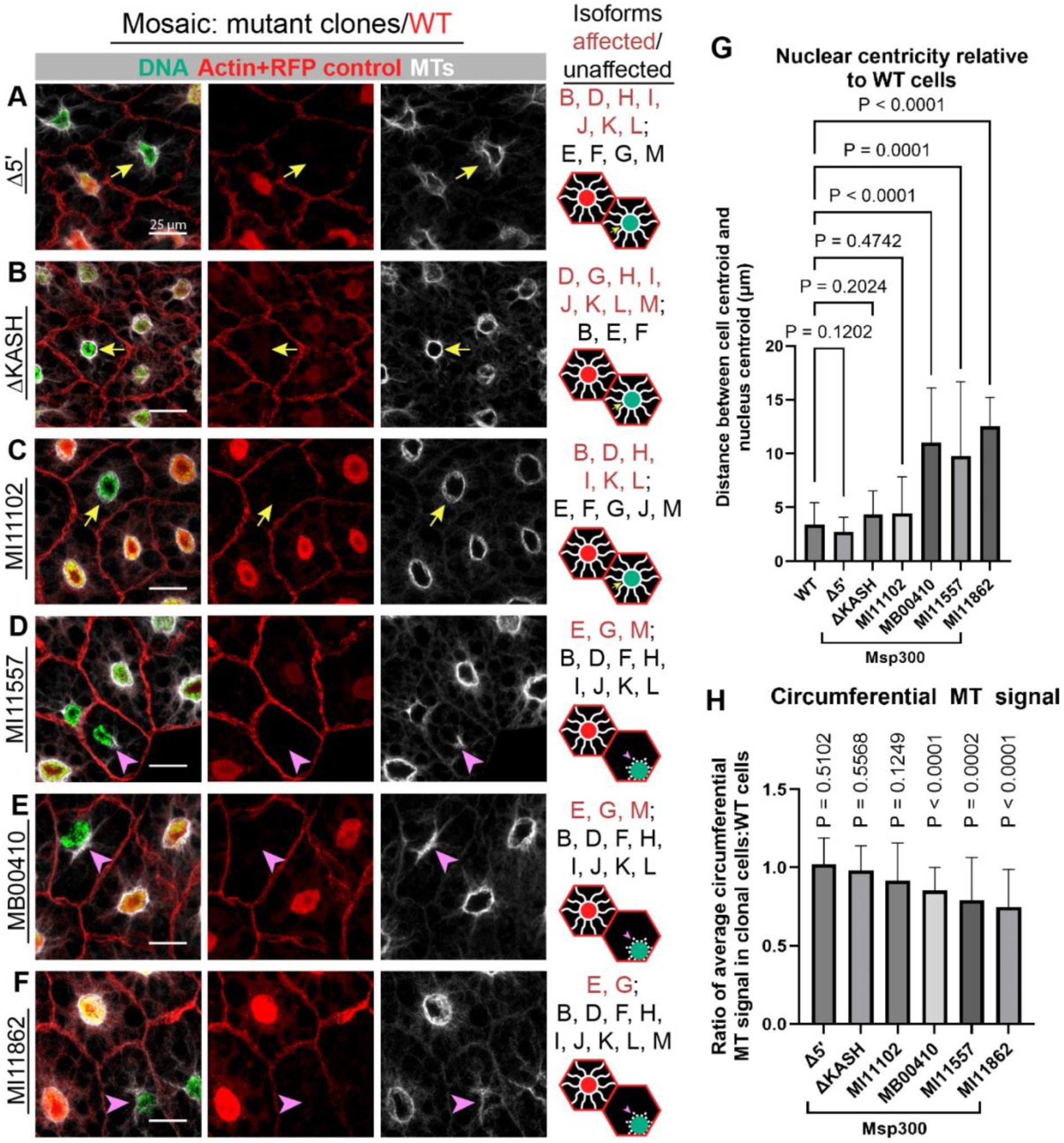
Msp300 isoforms E and G are required for nuclear positioning and microtubule organization. Nuclear positioning and MT organization was assessed by immunostaining fat bodies containing mosaic clones of Δ5′ (A), ΔKASH (B), MI11102 (C), MI11557 (D), MB00410 (E), and MI11862 (F) *Msp300* mutations. Representative images are shown. The FLP-FRT technique was used to generate mutant clones, where WT alleles are distinguished by the expression of nuclear RFP. The isoforms affected by each mutant allele are listed (red text). Normal nuclear positioning and MT organization is represented by a yellow arrow, while mispositioned nuclei with disrupted MT organization are indicated with a pink arrowhead. Disruption of isoforms essential to the ncMTOC results in the loss of nuclear centricity (G) and the reduction of circumferential MTs (H). Statistical significance was determined for nuclear centricity and circumferential MT signal using a one-way ANOVA with Dunnett’s T3 test and a one sample Student’s test with a theoretical mean of 1.000, respectively (21 ≤ *n* ≤ 30). Bars on the graph represent the mean ± SD of the data. Source data is present in S4 Data.

To probe the isoforms responsible for generating the ncMTOC further, we examined phenotypes for MiMIC transposable element insertion lines within *Msp300* exons that conceptually cause *Msp300* loss of function (see Fig 1A). Consistent with the 5′ deletion, the MI11102 [61] insertion within a coding exon near the 5′ end did not result in any changes to nuclear positioning or MT organization (Fig 3C, G-H), further supporting that all isoforms encoded by exons in the 5′ half of the gene are not required for generation of the ncMTOC. In contrast, disruption of isoforms E, G, and M by MI11557 [61] and MB00410 [62], or just E and G by MI11862 [61], results in severe nuclear mispositioning and disruption of radial and circumferential MTs (Fig 3D-F, G-H). These data show that these isoforms, or at least the coding regions downstream of these three transposon insertion lines, are necessary for proper function of the ncMTOC. In addition, the results show that disruption of isoforms E and G significantly impairs the fat body ncMTOC and that expression of isoform M is not sufficient to support the ncMTOC. We conclude from these mutant analyses that Msp300 isoforms E and/or G are required to generate the ncMTOC on the fat body nuclear surface.

In addition to targeting sets of isoforms with mutant lines, we also used RNAi to knock down sets of isoforms to complement and corroborate the mutant analysis described above, and also to further discern the isoforms required for the ncMTOC. We used the RNAi lines shown in Figures 1 and 2, and again generated mosaic knockdown with the coinFLP system, but in these experiments we assessed MT organization and nuclear positioning rather than Msp300 expression as done in Figure 2. First, we compared wild-type (*w^1118^*) to the internal control cells (Fig 4A, F-G). The HMS00368 RNAi line disrupted all Msp300 isoforms and resulted in severe nuclear mispositioning and MT disruption as shown previously [29] (Fig 4B, F-G). Knockdown of the isoforms expressed from the 5′ region of the gene did not affect nuclear positioning or MT organization (Fig 4C, F-G), consistent with the findings from mutants that affect the 5′ isoforms as shown in Figure 3. When targeting isoforms E and G with GD10406 RNAi, however, we observed mispositioned nuclei and disorganized MTs (Fig 4D, F-G). This result further shows that isoforms E and/or G are required to organize MTs at the ncMTOC. However, knockdown of isoform G and other isoforms containing the KASH domain using the GD10317 RNAi line resulted in moderate nuclear mispositioning and microtubule disruption (Fig 4E, F-G) indicating that isoform G supports the ncMTOC, but that isoform E is sufficient. We further show that when all isoforms except E and F are knocked down by RNAi (by expressing GD9539 and GD10317 together), the ncMTOC remains intact, whereas knockdown of all isoforms except F (GD9539 + GD10406) resulted in severe disruption of the ncMTOC, indicating that Msp300-PF is not sufficient (Fig S1). Together, the results from the mutants and the RNAi lines show that, while isoform G plays a contributing role in MT organization, isoform E is sufficient to generate the perinuclear ncMTOC. We were unable to test whether isoform G is also sufficient in the absence of E.

**Figure 4.**
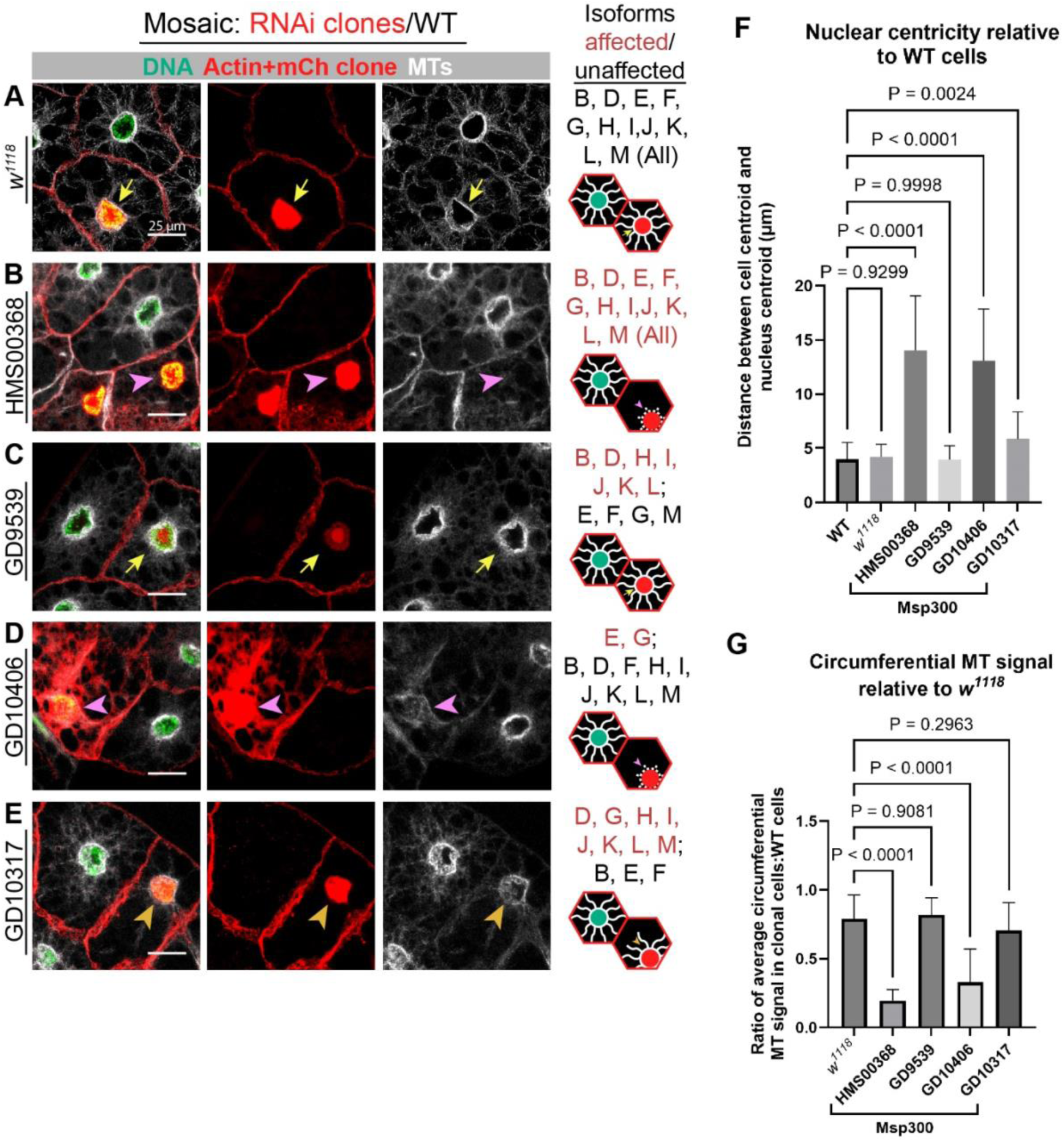
The perinuclear ncMTOC is organized by isoforms E and. **G.** Immunofluorescence microscopy images of WT (*w^1118^*) (A), HMS00368 (B), GD9539 (C), GD10406 (D), and GD10317 (E) RNAi lines to assess nuclear positioning and MT organization. Images shown are representative images. Mosaic clones were generated by the coinFLP technique, where knockdown clones are marked with nuclear mCherry (red). RNAi knockdown cells with properly positioned nuclei and organized radial and circumferential MTs are indicated with yellow arrows, and affected nuclei and MTs are indicated with pink arrowheads. Knockdown with GD10317 shows a moderate phenotype characterized by reduced MTs in the absence of nuclear mispositioning (gold arrowhead). Quantitation of nuclear centricity (F) and circumferential MT signal (G). For both analyses, statistics were conducted using a one-way ANOVA with Dunnett’s T3 test (*n* = 30). Bar graphs depict the mean ± SD. S4 Data contains the source data.

### Retrograde endocytic vesicle trafficking is dependent on Msp300 isoform E

In addition to nuclear positioning and MT organization, we assessed the effects of Msp300 isoform depletion on endosome trafficking. Prior work showed that Msp300 is required for the ncMTOC to organize MTs that support retrograde dynein-dependent endocytic vesicle trafficking [29]. In wild-type cells, Rab5 endosomal vesicles are at the highest density near the plasma membrane and perinuclearly, and disruption of MTs or dynein results in a loss of Rab5 endosome trafficking to the nuclear periphery [29]. To test the involvement of distinct Msp300 isoforms in endocytic vesicle trafficking, we examined the distribution of the endosomal marker GFP-Rab5 upon disruption of select isoform(s) by RNAi knockdown in the larval fat body using SPARC-GAL4. In the *w^1118^* control cells and cells expressing *Luciferase^JF01355^* (RNAi control), Rab5 vesicles comprise a perinuclear population forming a halo-like pattern (Fig 5A-B, H). Knockdown of *αTub84B* (positive control) causes impaired retrograde trafficking, as seen by the loss of the perinuclear Rab5 signal (Fig 5C, H). Similarly, the knockdown of all *Msp300* isoforms using HMS00368 RNAi results in diminished Rab5 vesicles at the nuclear surface (Fig 5D, H). Disruption of the 5′ isoforms by the GD9539 RNAi line did not impact the Rab5 distribution pattern, indicative of normal endosome trafficking (Fig 5E, H). In contrast, knockdown of isoforms E and G with GD10406 RNAi resulted in reduced perinuclear GFP-Rab5 endosome signal (Fig 5F, H). The knockdown of Isoform G and other KASH-containing isoforms via GD10317 RNAi did not appreciably reduce the perinuclear GFP-Rab5 signal (Fig 5G, H). Together, these results further show that isoform E is sufficient for ncMTOC function by supporting endosomal vesicle trafficking in the fat body.

**Figure 5.**
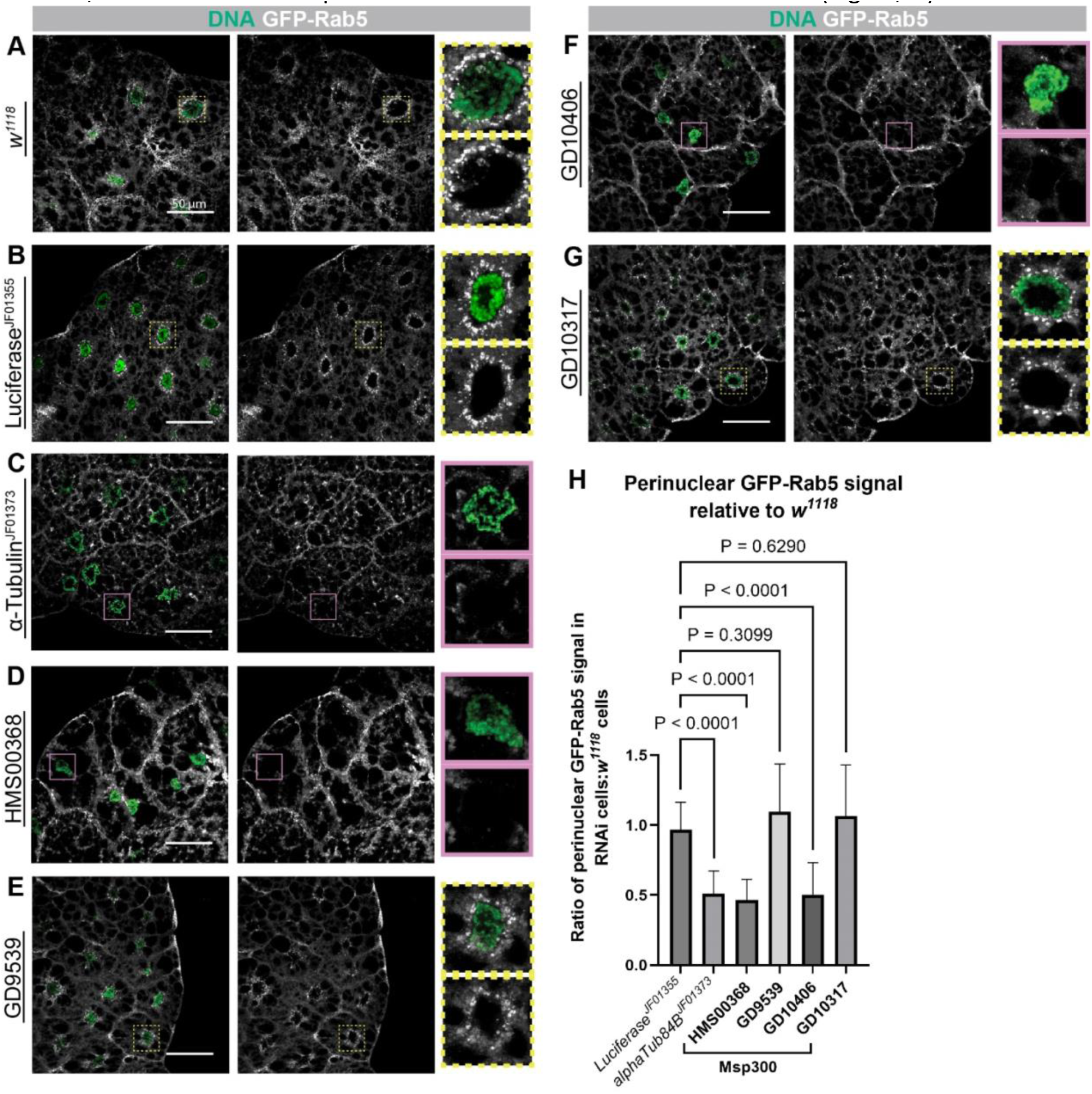
Msp300 isoforms E+G are required for retrograde trafficking of endocytic vesicles. Analysis of retrograde endosome trafficking in *w^1118^*(A) cells, *Luciferase^JF01355^* (B), and *alphaTub84B^JF01373^*(C), as well as in cells expressing the HMS00368 (D), GD9539 (E), GD10406 (F), and GD10317 (G) RNAi lines. Representative images are shown here. RNAi was expressed throughout the entire fat body tissue with SPARC-GAL4. Images were captured using the same parameters to allow for direct comparison between samples. Insets show a magnified view of a single nucleus (green) and the perinuclear population of Rab5 endosomes (white). Control or unaffected cells exhibit a halo-like pattern of endocytic vesicles surrounding the nucleus (yellow dashed boxes), while knockdown of Msp300 isoforms essential for the MTOC causes loss of this signal (pink boxes). (H) Bar graph depicting perinuclear GFP-Rab5 signal. A one-way ANOVA with Dunnett’s T3 test was used for statistical analysis (*n* = 30). The mean ± SD is shown in the graph. Source data is present in S4 Data.

### Msp300 isoform E is required for the recruitment of the ncMTOC components shot and Patronin

To further evaluate the role of Msp300 isoforms at the fat body ncMTOC, we examined their involvement in the recruitment of other ncMTOC components. Msp300 recruits the spectraplakin short stop (shot) and the microtubule minus-end stabilizer Patronin to the ncMTOC [29, 30]. In the *w^1118^* control, shot localizes to the nuclear surface, along MTs, and on the plasma membrane (Fig 6A, F). When all *Msp300* isoforms were disrupted, shot no longer localizes to the nuclear surface [29]. Consistently, we observed a decreased amount of shot at the nuclear surface when we used HMS00368 RNAi to knock down all isoforms (Fig 6B, F). The knockdown does not, however, affect the localization of shot at the plasma membrane. Interestingly, knockdown of the 5′ isoforms (with GD9539 RNAi) resulted in an increase in the perinuclear shot signal (Fig 6C, F). Knockdown of the KASH isoforms that include Isoform G also showed an increase in the perinuclear localization of shot (Fig 6E, F). Targeting of isoforms E and G with GD10406, on the other hand, causes a loss of shot from the nuclear surface (Fig 6D, F). These results indicate that Msp300 isoform E is sufficient for the recruitment of shot to the ncMTOC, consistent with its role in establishing the perinuclear ncMTOC.

**Figure 6.**
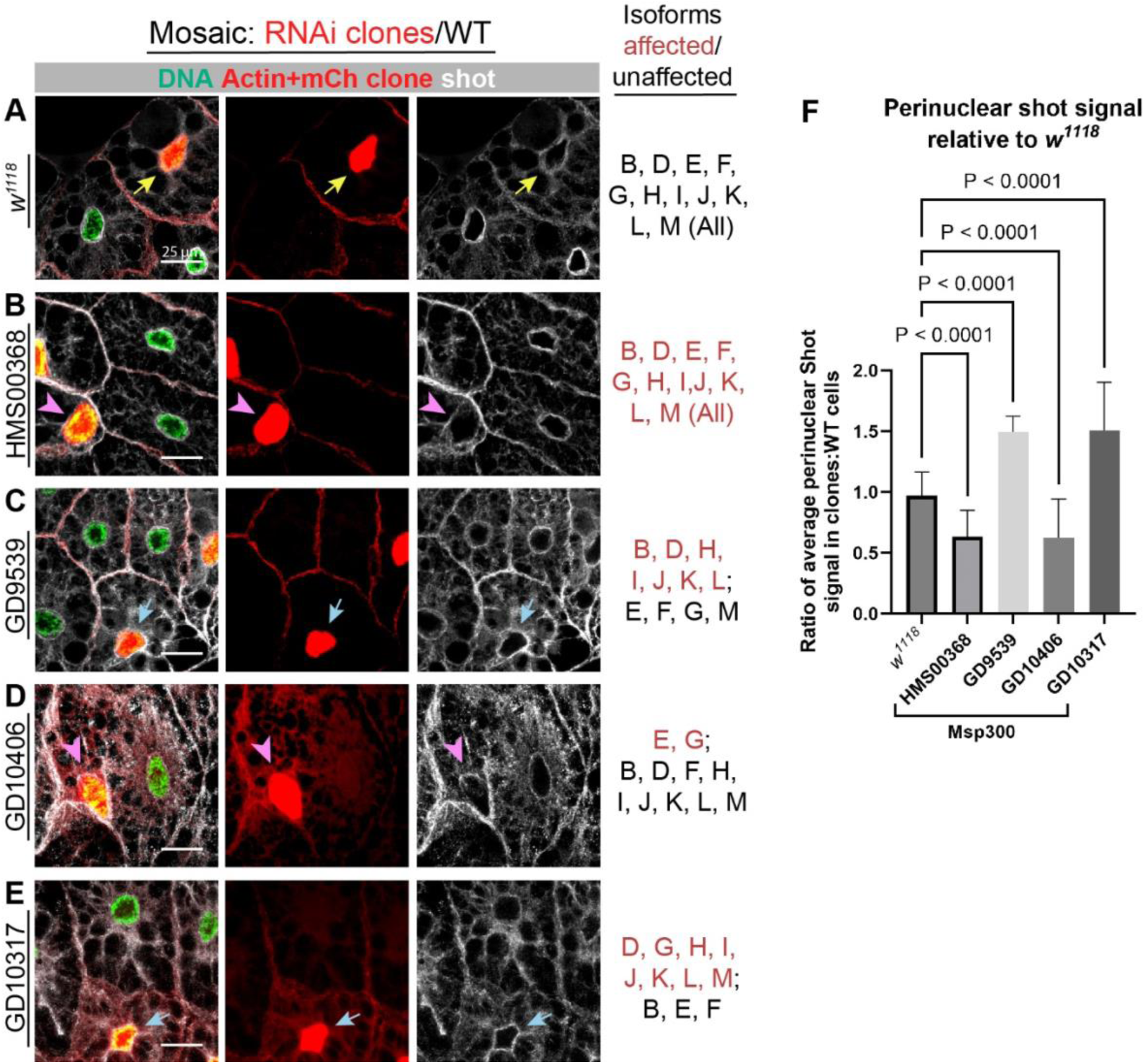
Shot localization to the ncMTOC requires isoform E. Fat bodies were stained for shot in mosaic tissues expressing *w^1118^* (A), HMS00368 (B), GD9539 (C), GD10406 (D), or GD10317 (E). Images shown are representative. The coinFLP system was used to express the RNAi, marked with co-expression of nuclear mCherry (red). Recruitment of shot to the nuclear surface by Msp300 results in a perinuclear shot signal (yellow arrows), while the knockdown of necessary Msp300 isoforms causes the loss of this signal (pink arrowheads). (F) Quantitation of perinuclear shot signal. Statistical analysis was performed via a one-way ANOVA and Dunnett’s T3 test (*n* = 30). Data are depicted as mean ± SD. S4 Data contains the source data.

Finally, we investigated the contribution of the isoforms to the recruitment of the CAMSAP ortholog Patronin to the ncMTOC. Patronin is a MT minus-end stabilizer and a major regulator of MTs at the fat body perinuclear ncMTOC and complete *Msp300* knockdown results in the loss of Patronin localization at the nuclear surface [29]. In *w^1118^*control and *Luciferase^JF01355^* control RNAi cells expressing Patronin-GFP, Patronin localizes to the nuclear surface (Fig 7A-B, H), while in *Patronin^HMS01547^* RNAi cells perinuclear Patronin is significantly reduced (Fig 7C, H). Disruption of all isoforms (with HMS00368 RNAi) results in a significant decrease in the perinuclear localization of Patronin (Fig 7D, H). Knockdown of the 5′ isoforms, in contrast, does not affect the presence of Patronin at the nuclear surface (Fig 7E, H). Patronin perinuclear localization is diminished when Isoforms E and G are knocked down with GD10406 RNAi but not when the 3′ KASH-containing isoforms are knocked down (Fig 7F-G, H), consistent with the role for isoform E in MT organization and shot recruitment to the nuclear surface. Together, these data further show that Msp300 isoform E is necessary and may be sufficient for MTOC generation through the recruitment of shot and Patronin.

**Figure 7.**
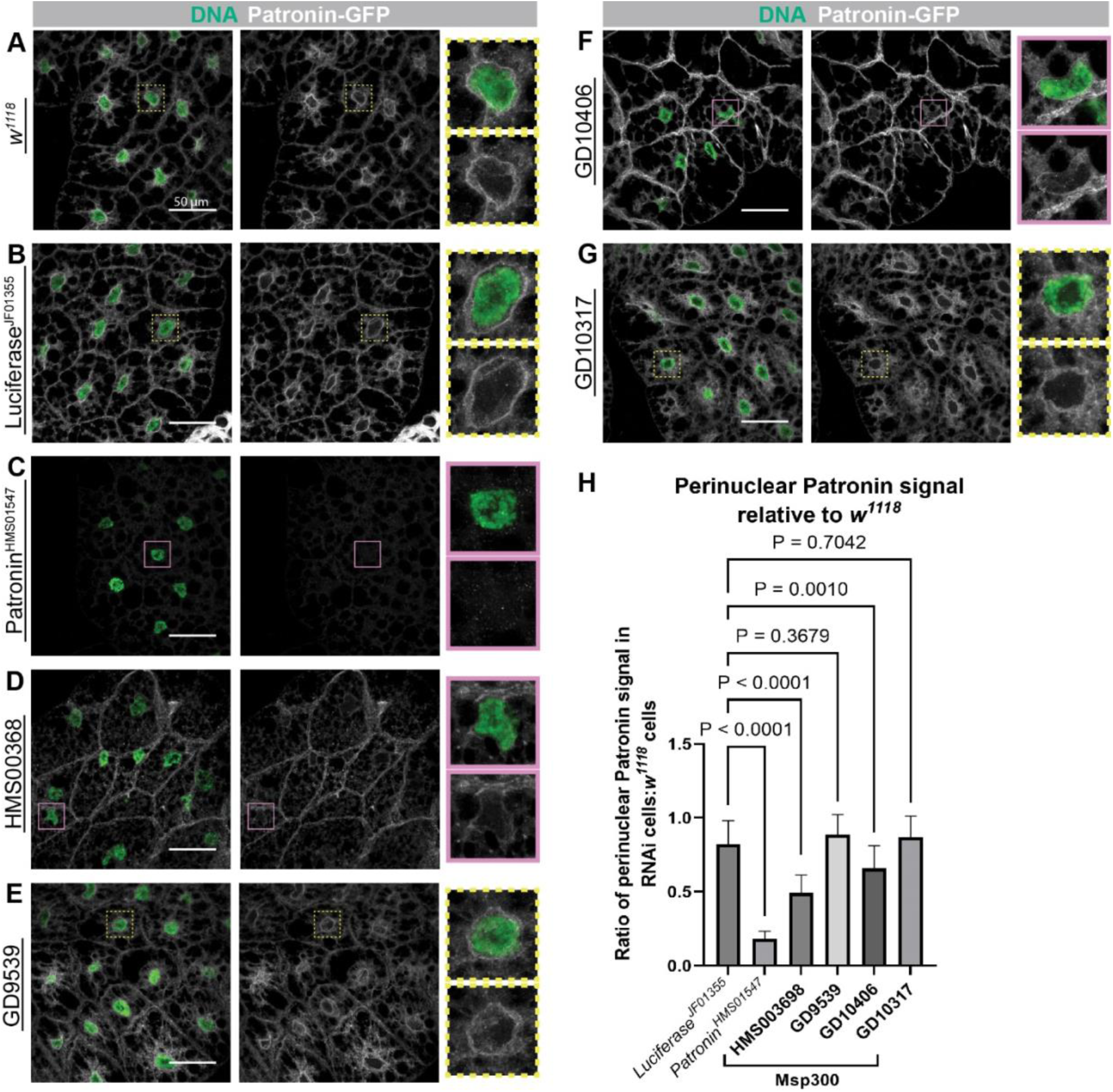
Patronin localization to the ncMTOC requires isoform E. Patronin localization in *w^1118^* (A), *Luciferase^JF01355^* (RNAi) (B), and *Patronin^HMS01547^* (RNAi) (C) fat bodies and fat bodies expressing the *Msp300* HMS00368 (D), GD9539 (E), GD10406 (F), or GD10317 (G) RNAi lines. Representative images are shown. RNAi lines were expressed throughout the fat body with SPARC-GAL4. The same confocal microscope parameters were used to capture the images for each sample. Nuclei (DAPI, green) and Patronin-GFP (white). Patronin normally localizes perinuclearly (yellow dashed boxes). Knockdown of Msp300 isoforms essential for the ncMTOC results in loss of Patronin signal at the nuclear surface (pink boxes). (H) Perinuclear Patronin signal in each sample represented in the form of a bar graph. Data is presented as the mean ± SD. Statistics were conducted via a one-way ANOVA with Dunnett’s T3 test (*n* = 30). Source data is present in S4 Data.

### Msp300-PE has two novel large tandem repeats and a KASH-like C-terminal domain

Most cell types do not generate an MTOC on the nuclear surface [63], and fat body cells appear to utilize a special isoform of Msp300, Msp300-PE to assemble it. To further understand what makes Msp300-PE unique, we analyzed its sequence and predicted domain structure with *in silico* analysis tools. Msp300-PE contained no other identified domains besides the SRs and the transmembrane domain based on analysis via the PredictProtein tool [64]. Intriguingly, Msp300-PE encodes two unique types of novel tandem repeat regions (Fig 8A-D), identified using the XSTREAM algorithm [65]. The first type, “Type 1,” consists of ∼20 copies of a 108-amino acid sequence that is highly similar among the tandem repeats. “Type 2” repeats consist of 101 amino acids tandemly repeated ∼5-7 times in three clusters (labeled 2A-2C in Figure 8) with a total of 19 repeats. These repeat domains have no previously described function. Using AlphaFold 3, we analyzed the predicted structures of the repeats [66]. Individually, each cluster of repeats is predicted to form similar structures despite no sequence similarity, comprised of beta strands organized in parallel and antiparallel, to form a folded-back sandwich structure with disordered loops present throughout the protein (Figure 8E-H, and Supplemental Videos 1 and 2). AlphaFold 3 predicts a loose association between the repeats that might induce further flattening of the structures (Fig 8I). In addition, the predicted structure of the C-terminus of Msp300-PE is shown. Similar to the KASH domain of other Msp300 isoforms (Fig 8J), the C-terminal domain (CTD) of Msp300-PE is predicted to contain a membrane-spanning α-helix and a mostly disordered CTD predicted to reside within the intermembrane space (Fig 8K).

**Figure 8.**
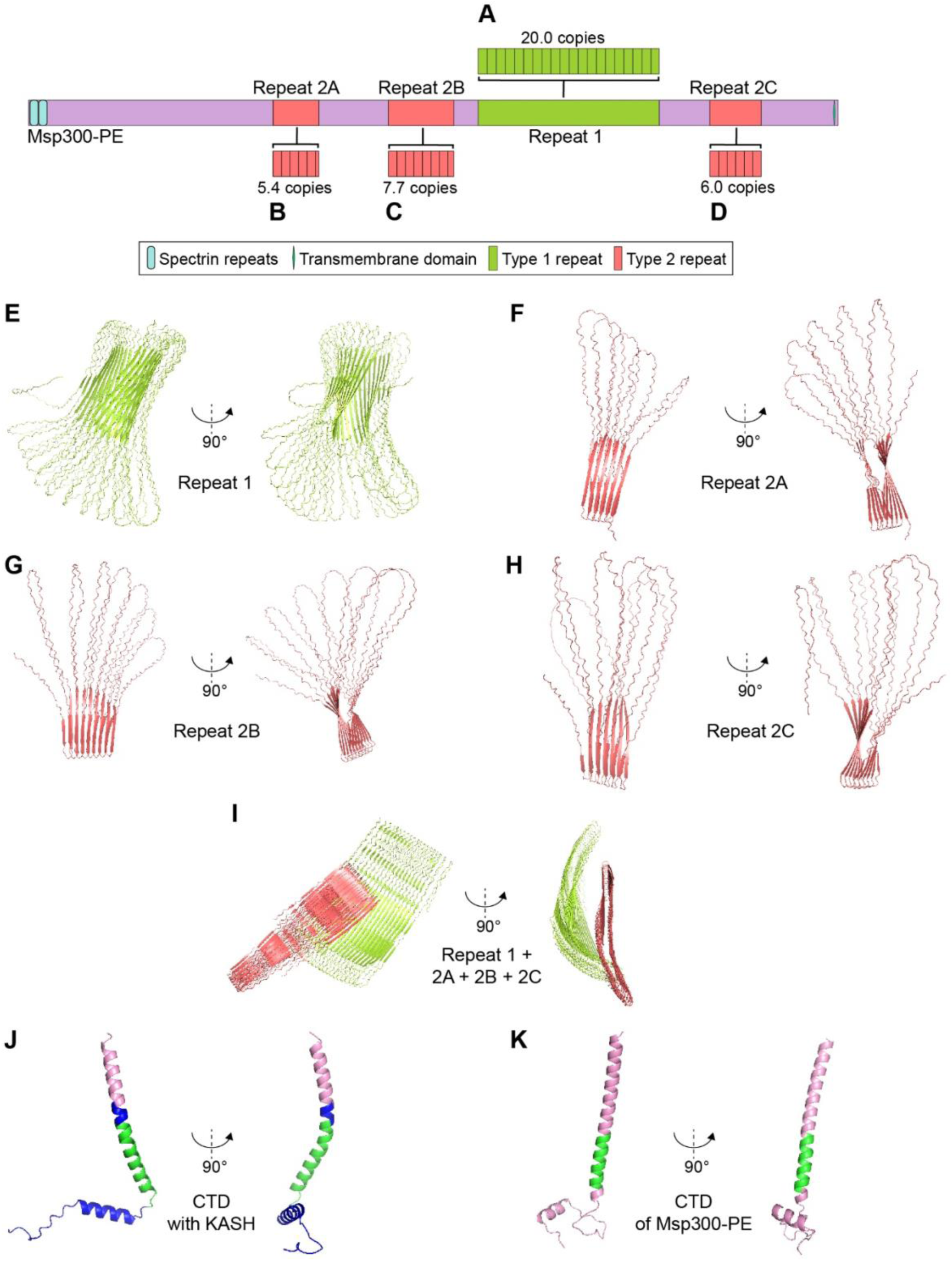
Msp300-PE contains two novel large tandem repeats and a KASH-like C-terminal domain. (A) Repeat 1 (lime green) consists of a 108-amino acid sequence repeated 20 times. (B-D) Repeat 2 (salmon) consists of a different 101-amino acid sequence repeated at least 19 times among 3 clusters. (E) Predicted structure of Repeat 1 and view of Repeat 1 rotated 90° about the *y*-axis. (F) Repeat 2A predicted structure before rotating 90° around the *y*-axis and after. (G) Predicted structure of Repeat 2B and view after rotating 90°C about the *y*-axis. (H) Repeat 2C predicted structure and view of Repeat 2C following 90° rotation around the *y*-axis. (I) Predicted structure of Repeat 1 (lime green) and Repeat 2A-2C (salmon) before and after 90° rotation about the *y*-axis. (J) Predicted structure of the CTD containg the KASH domain (blue), transmembrane domain (green), and linker region (lavender) and view after 90° rotation around the *y*-axis. (K) Predicted structure of the CTD of Msp300-PE with the transmembrane domain (green) and linker region (lavender) shown before 90° rotation around the *y*-axis and after.

## Discussion

Here we show that a unique isoform of Msp300, Msp300-PE, is necessary in fat body cells to organize the ncMTOC on the nuclear surface. Msp300-PE and the isoforms expressed from the same transcriptional promotor (Msp300-PG and -PM) are expressed most highly in the fat body compared to the expression of the other isoforms, indicating that these isoforms and the unique protein domains they encode may be responsible for assembly of the ncMTOC on the nuclear surface. Our data also indicate that Msp300-PE is sufficient to generate the ncMTOC on the nuclear surface. Msp300-PE organizes the microtubules through recruitment of shot and Patronin, a major regulator of microtubules at the perinuclear ncMTOC [29] and is consequently required for nuclear positioning and endocytic vesicle trafficking in fat body cells. These findings establish why Msp300, expressed from a complex locus with expression in many cell types, organizes an ncMTOC on the surface of nuclei in a cell-type specific manner. Msp300-PE has a unique domain structure, containing a predicted novel outer nuclear membrane targeting domain and with novel repeat domains which, combined, indicate that Msp300-PE is a novel alternative LINC complex component.

The knockdown of Msp300-PE/PG, or their disruption together with Msp300-PM with transposon insertions, indicates that Msp300-PE and -PG organize the perinuclear MTOC in fat bodies. The loss of Msp300-PG, together with all KASH-containing isoforms, altered nuclear positioning but did not significantly block the function of the MTOC, as MTs and the localization of Rab5-GFP were not significantly affected. Thus, we infer from all the results that Msp300-PE is sufficient to organize the MTOC, while Msp300-PG contributes. It is possible that Msp300-PG is also sufficient, but we did not have the genetic tools to test that in this study. Due to the technical limitations of expressing such large constructs, we were unable to test sufficiency by expressing a transgene of Msp300-PE/PG in a null background; the Msp300-PE coding sequence is almost 29,000 bp in length and Msp300-PG is longer (over 40kbp). Furthermore, while the existence of PE/PG is supported by the available cDNAs corresponding to the exonic regions common to both isoforms, further validation is required to confirm the existence of other isoforms such as PF [48].

In contrast with the majority of the other Msp300 isoforms, and nesprins generally across species, isoform E lacks the characteristic KASH domain, the CH domains, and the high number of spectrin repeats. Also, we show that loss of all KASH-containing isoforms does not affect the ncMTOC in the fat body or the localization of Msp300 to the nuclear surface. However, KASH-less isoforms have been identified for other nesprins in Drosophila and other species including *Drosophila* klar, vertebrate nesprin-1/2, and *C. elegans* ANC-1. A KASH-less isoform of klar was shown to be associated with lipid droplets [67, 68]. In addition, KASH-less splice variants of nesprin-1 and nesprin-2 have been characterized [69–71], and an isoform of nesprin-1 lacking the KASH domain is proposed to be expressed in the embryonic mouse brain [72]. A nesprin-1-giant isoform lacking the KASH domain has been identified in the CNS and is associated with spinocerebellar ataxia [73, 74]. Moreover, In *C. elegans*, ANC-1 regulates nuclear positioning in hypodermis syncytia independently of the KASH domain and the LINC complex [75]. Furthermore, while in *Drosophila* loss of the 5′ isoforms was lethal at the first instar larval stage, deletion of the KASH domain from all isoforms was viable [59]. In Drosophila muscle, however, Msp300 is required for nuclear positioning in a KASH-dependent manner [21].

Despite the similarities with other KASH-less isoforms, isoform E is particularly unique due to the presence of the two types of highly conserved ∼100 amino acid-long tandem repeats. Based on the results of a protein BLAST against all insects, the repeats appear to possess significant homology only among Drosophila species. In addition, the repeats are characterized by an overall negative charge, as observed in intrinsically disordered regions of proteins [76, 77]. The negatively charged regions may also facilitate binding with other proteins. For example, the positively charged regions of Patronin, which bind to the negatively charged regions of MTs, may also interact with the repeats of Msp300. The unique domain structure of Msp300-PE is likely linked to the mechanism by which this variant nesprin organizes the ncMTOC. Furthermore, the structural similarity of the C-terminal domain of Msp300-PE with the KASH domain of other Msp300 isoforms may provide a binding site for a previously unidentified inner nuclear membrane protein partner of Msp300-PE. Together, Msp300-PE and the predicted inner nuclear membrane protein may form a novel LINC complex.

Overall, this work shows that Msp300-PE has unique function to generate a perinuclear ncMTOC. Future work will elucidate the functions of the unique domains and how Msp300-PE is able to generate an ncMTOC on the surface of nuclei.

## Materials and Methods

### Genetic data

Information pertaining to the gene sequence and isoforms was obtained from the data available on FlyBase (https://flybase.org/reports/FBgn0261836.htm) [45].

#### Drosophila stocks

Information for each *Drosophila* stock is contained in Supplementary Table 1. Flies were raised using standard food made of cornmeal, molasses, and yeast and were maintained at 25°C.

#### Generation and expression of FLP-FRT and coinFLP stocks

The generation of FLP-FRT mutant clones was conducted by crossing *y^1^ w* hsFLP^22^*; Ubi-RFP-nls neoFRT40A females to males with each *Msp300* allele recombined with neoFRT40A. For each cross, 30-40 females were used.

The generation of coinFLP mosaic RNAi knockdowns was conducted using an adaptation of the coinFLP system [55]. For each cross, 25-35 females of the genotype *w* hsFLP^1^ coinFLP-GAL4^attP3^*;; UAS-mCherry-nls3 were crossed to RNAi line males.

Crosses were established for at least 2-3 days before proceeding with heat shock to induce recombination. FLP-FRT and coinFLP crossed flies were transferred to fresh food vials for 1-2 hours, then to fresh food vials to lay eggs for 2-3 h, and then adults were removed and embryos were aged past the cleavage stage at 29°C for 2 hr. Embryos were then heat-shocked in a 37°C water bath for 30 minutes and then returned to 29°C for coinFLP crosses or 25°C for FLP-FRT crosses. Wandering stage larvae were collected 4-5 days later for fat body dissection.

#### Sample preparation and immunostaining

Fat bodies from late third-instar wandering larvae were dissected in 1× Dulbecco’s phosphate-buffered saline (D-PBS) (GIBCO, Waltham, MA, USA) as previously described [29]. Fat bodies from two larvae were used for each slide. Following dissection, samples were transferred to a poly-L-lysine-coated slide. For Patronin-GFP samples, the fat bodies were fixed in 15 µL of 100% methanol under a siliconized coverslip for 10 min at −20°C. The coverslip was removed and the slides were placed in 1× PBS (137 mM NaCl, 2.7 mM KCl, 8.0 mM Na_2_HPO_4_, and 1.4 mM KH_2_PO_4_). All other samples were fixed in 15 µL of 4% paraformaldehyde (PFA) for 8 min before a siliconized coverslip was placed on the slide. The sample was gently flattened and then flash frozen in liquid nitrogen. While still frozen, the coverslip was removed and the slides placed in 1× PBS. For all samples, a ∼2 cm diameter hydrophobic ring was drawn around the sample with a Super Pap Pen (Electron Microscopy Sciences, Cat # 71312). Fat bodies were permeabilized, blocked, and stained in 100 µL of staining solution (1× PBS, 1% BSA, 0.1% saponin) for 2 hr at RT or overnight at 4°C using the antibodies and dyes listed in Supplementary Table 2. Slides were extensively washed in 1× PBS following staining. Samples were mounted in 10 µL of mounting medium (80% glycerol, 0.1 M Tris pH 8.8, 0.05% p-phenylenediamine dihydrochloride (Sigma, cat. no. P1519)) and stored at −20°C.

#### Image acquisition

Fat bodies were imaged with either a Nikon A1 or Nikon AX laser scanning confocal microscope (Nikon, Japan). Specimens were imaged using a 60× oil-immersion objective with a 1.49 numerical aperture (NA). Images were captured with the Nikon NIS-Elements AR software (version 4.6 or 5.42 with Nikon A1 or Nikon AX, respectively) with 0.5 µm between *z*-sections. All images are presented as maximum intensity projections of the *z*-stack.

#### Quantification analysis

Microscopy images were quantified using Fiji [78]. Perinuclear localization was calculated from a single *z*-slice, while nuclear mispositioning was calculated from a maximum intensity projection. Quantification of perinuclear localization was performed by creating multiple ROIs of the same size throughout the perinuclear region and averaging the total fluorescence intensity of each ROI. The background for each cell was accounted for by subtracting the value of an ROI in the cytoplasm from the value of the perinuclear region. For samples utilizing the coinFLP system, the value for each clone was depicted as ratio of the clone to an adjacent wild-type control cell. For all other samples, the average value of the clones from a slide was divided by the average value of the corresponding *w^1118^* control slide. Quantification of nuclear positioning was conducted as previously described [29]. In brief, the coordinates of the cell or nuclear centroid were obtained by defining the boundary of the plasma membrane or nucleus, respectively. Next, the distance between the two coordinates was calculated.

#### RT-qPCR

For preparation of whole fly and flight muscle RNA, 10 adult flies were used. Ovary RNA was extracted from 10 adult virgin female flies. Brains and fat bodies were dissected from 10 late third-instar wandering larvae. Tissues were washed in 1× D-PBS following dissection. After homogenizing sample via pellet pestle and aspiration with a 20 G needle, RNA was isolated from all samples using TRlzol (Ambion, Austin, TX, USA) following the recommended protocol from the manufacturer. For all extraction steps RNase-free/DNase-free plasticware and solutions were used. Isolated RNA samples were resuspended in UltraPure DNase/RNase-free distilled water and treated with TURBO DNA-*free* kit (Thermo Fisher, Waltham, MA, USA). cDNA was produced using the SuperScript IV (Invitrogen, Waltham, MA, USA) kit with random hexamers primers. Quantitative PCR was performed using the primer pairs shown in Figure 1A along with those for the *αTub84B* [79] and *Gpdh1* reference genes. Primer sequences are listed in Supplementary Table 3. Amplification and quantification were performed using the Brilliant III Ultra-Fast qPCR Master Mix (Agilent, Santa Clara, CA, USA) at a Bio-Rad CFX384 real-time analyzer. Quantification was performed using the 2^-ΔΔCt^ method. The expression levels of the 5′ and E, G, M isoforms were compared to the average expression levels of All isoforms amplified from two separate primer pairs.

#### Statistics and reproducibility

For microscopy image calculations, three slides, with each containing the fat bodies from two larvae, were imaged. On each slide, ≥ 5 wild-type and ≥ 5 clone cells were included in the quantitation. The use of the mosaic systems in this study provided an internal control, which reduces variability arising from sample preparation. For non-mosaic tissues, 10 cells from each slide were quantified. Nuclear positioning and perinuclear localization values are expressed as the mean ± standard deviation (SD). All samples for RT-qPCR were dissected in triplicate and each reaction was performed in triplicate. The data were significant based on a *P*-value < 0.05. For datasets with negative control samples, a one-way ANOVA was performed, while a one sample *t*-test was used for all other datasets. GraphPad Prism 10 (Dotmatics, Boston, MA, USA) was used for all analyses.

#### In silico analysis

Analysis of Msp300-PE was performed using the publicly available amino acid sequence (NCBI Reference Sequence: NP_001188693.1). PredictProtein [64] was used to identify the residues predicted to comprise the transmembrane domain of Msp300-PE. The locations of the tandem repeats were determined via XSTREAM [65]. Structure predictions of the repeat regions and C-termini of PD and PE were performed with AlphaFold 3 [66]. Predicted structures were visualized using PyMOL (The PyMOL Molecular Graphics System, Version 2.4.0 Schrödinger, LLC). All analyses were conducted using the default parameters of the tools.

#### Graphics

All graphics were made using Adobe Illustrator (version 25.3.1). Also, all figures were composed using Adobe Illustrator.

## Acknowledgements

We thank Véronique Morel, Bénédicte Durand, Talila Volk, Michael Welte, and the Bloomington *Drosophila* Stock Center for *Drosophila* stocks; Talila Volk and the Developmental Studies Hybridoma Bank for antibodies; and Yue Julia Wang for assistance with RT-qPCR. We are grateful to Bloomington Drosophila Stock Center (NIH P40OD018537), Vienna Drosophila Resource Center [57], and FlyBase [45] for curating indispensable tools and resources. Thank you to Batory Foods for their generous donation of fly food reagents to support this work.

This work was supported by NIH grant R01GM139971 to T. Megraw, and National Natural Science Foundation of China grant (32370731) and Shenzhen Science and Technology Program grant (JCYJ20230807091308018) to Y. Zheng.

