## Supplemental Figure 1 for "A distinct isoform of Msp300 (nesprin) organizes the perinuclear microtubule organizing center in adipocytes"

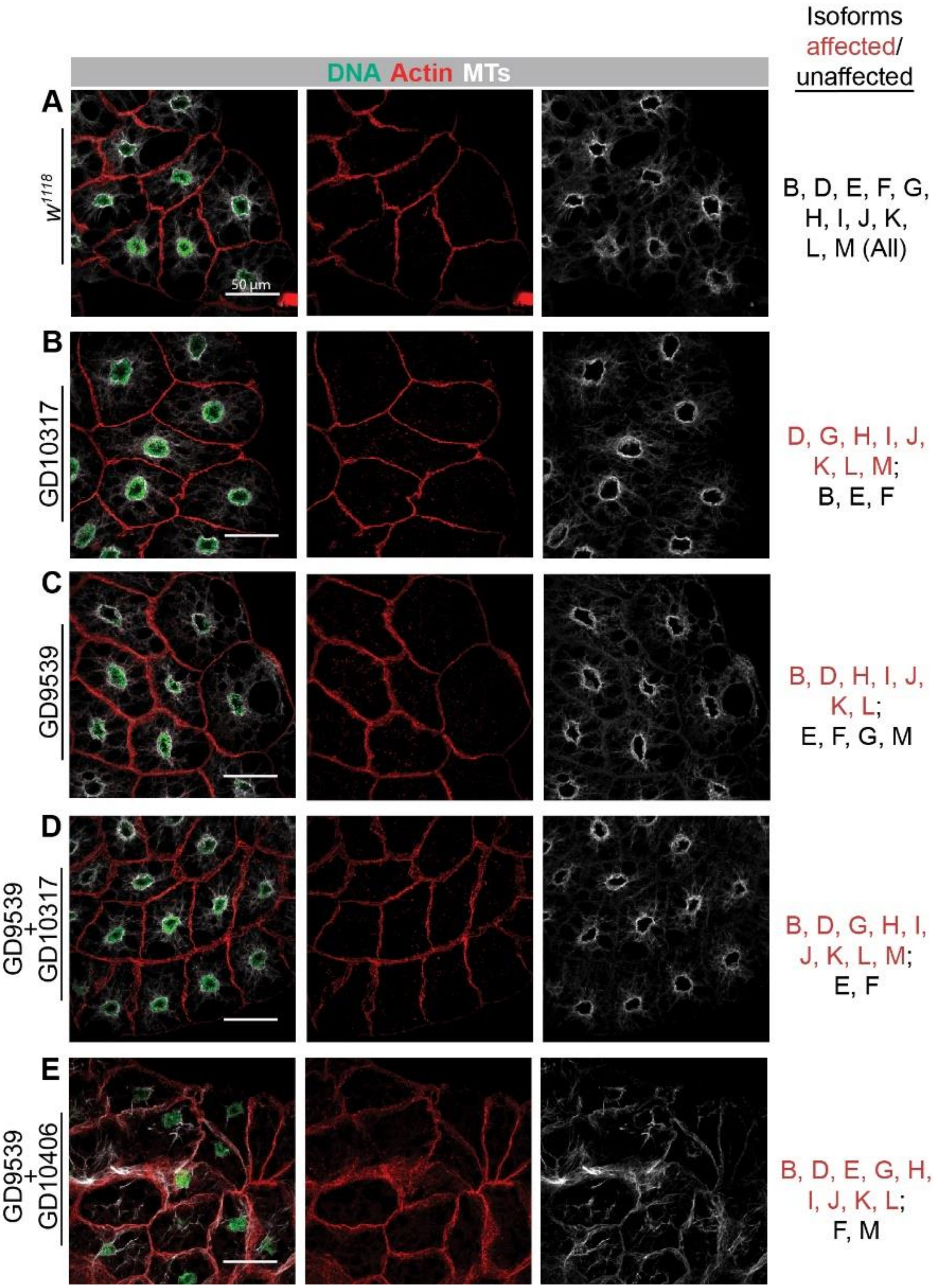


**Figure S1. Isoform E is necessary and sufficient to organize the ncMTOC.** Analysis of nuclear positioning and microtubule organization in $w^{1118}$ (A), GD10317 (B), GD9539 (C), GD9539+GD10317 (D), or GD9539+GD10406 (E) RNAi lines expressed with SPARC-GAL4. All samples have SPARC-GAL4, including "$w^{1118}$", which is the progeny of SPARC-GAL4 crossed to $w^{1118}$. The same imaging parameters were used for all samples. The isoforms knocked down (affected) and unaffected by each RNAi line are listed.

### Supplemental Video captions

**Supplemental Video 1. Predicted structure of Repeat 1.** Video showing the structure of Repeat 1 as predicted by AlphaFold 3. The structure is rotated around the *y*-axis. Movie made with PyMOL.

**Supplemental Video 2. Predicted structure of Repeat 2A.** Video showing the structure of Repeat 2A as predicted by AlphaFold 3. The structure is rotated around the *y*-axis. Movie made with PyMOL.

**Supplemental Video 3. Predicted structure of the interaction between Repeat 1 and Repeat 2A-C.** Video showing the structures of Repeat 1 (lime green) and Repeat 2A-C (salmon) as predicted by AlphaFold 3. The overall structure is rotated around the *y*-axis. Movie made with PyMOL.
